# Evaluation of Open Hollow Hydroxyapatite Microsphere on Bone Regeneration in Rat Calvarial Defects

**DOI:** 10.1101/669598

**Authors:** Youqu Shen, Mohamed Rahaman, Yongxian Liu, Yue-Wern Huang

## Abstract

Hollow hydroxyapatite (HA) microspheres showed the ability to facilitate bone regeneration in rats with non-healing calvarial defects. However, new bone formation in the rat calvarial defect implanted with the closed HA microspheres was limited. The objective of this work is to evaluate size-, time, and structure-dependent bone regeneration between open and closed HA microspheres in an osseous model. Open HA microspheres were obtained by sectioning closed HA microspheres. The open HA microsphere had dense convex surface and rough and porous concave surface. For both size ranges (ϕ106-150 μm vs. ϕ212-250 μm), the open HA microsphere were more effective in facilitating bone regeneration than the closed HA microspheres in rat calvarial defects. Bone regeneration in the open HA microspheres (49 ± 7% for ϕ106-150 μm; 40 ± 8% for ϕ212-250 μm) were higher than the closed HA microsphere (26 ± 8% for ϕ106-150 μm; 30 ± 9% for ϕ212-250 μm) at 12 weeks. Furthermore, the open HA microspheres of smaller size showed a significant increase in bone regeneration than the open HA microspheres of larger size at both 6 weeks and 12 weeks. The difference in bone regeneration between these microspheres could be due to their differences in microstructures, namely curvature, concavity, porosity, surface roughness, and total surface area available for cells to attached to.

## Introduction

Effective regeneration of bone defects caused by trauma or chronic diseases is a significant clinical challenge. Over the past few decades, researchers have investigated the mechanism of bone regeneration to better inform the designs of healing strategies [1–3]. Bone healing involves three primary stages: the early inflammatory stage; the repair stage and the late remodeling stage [4]. These three stages are distinct, but continuous. In the inflammatory stage, a hematoma forms and inflammatory cells infiltrate the bone, resulting in the formation of granulation tissue, vascular tissue and immature tissue. During the repair stage, new blood vessels are developed to facilitate tissue regeneration and a soft callus is formed around the repair site. Bone healing is completed during the remodeling stage in which the bone is restored to its original shape, structure and mechanical strength.

Clinically, bone deficiency is overcome using treatments that rely on bone regeneration and augmentation. While various treatments have been investigated with encouraging results [5], complete and predictable bone reconstruction is often difficult [6]. Autologous bone grafts are the gold standard for treatment because they contain osteoinductive growth factors, osteogenic cells and a structural scaffold. However, disadvantages of this treatment include limited tissue availability, increased surgery time, additional pain and cosmetic imperfection at the donor site [6–8]. Many of these issues can increase the health care cost for the patient [9]. An alternative to autogenous bone is allogenic bone, which can induce moderate healing results due to its preserved osteoinductivity. However, allografts are costly, can have unpredictable effects on growth due to donor variance, cause adverse immune reactions, and increase the risk of disease transference [10–12]. Synthetic bone grafts have advantages such as consistent quality, safety, and good tissue tolerance, but they usually function as inert or merely osteoconductive implants. Encouraging results have been reported.

Hydroxyapatite (HA), the main component and essential ingredient of human bone, can be prepared by chemical reactions. Studies have demonstrated that HA supports bone regeneration and bonding to surrounding tissue because of its biocompatibility, bioactivity, and osteoconductivity [13]. Our studies with the closed HA microspheres showed the ability to regenerate bone in non-healing rat calvarial defects [14, 15]. Experiments with ϕ106-150 𝜇m and ϕ150-250 𝜇m closed HA microspheres showed differences in mechanical properties and biological tests [16]. The size variation of closed HA microspheres could affect the structure of HA microspheres. The changes in structure can influence on the biological tests in return. We sporadically observed that there tended to be better bone regeneration with broken closed microspheres with micro-concavity [15, 17]. This observation motivated us to design this study that focused on enhanced bone regeneration with open microspheres. We hypothesize that open HA microsphere with special geometric characters can yield better bone regeneration compared with the closed HA microspheres. Our goal is to investigate whether bone regeneration in an osseous model is microgeometry-, size-, and time-dependent. To achieve our goal, two size ranges (ϕ106-150 μm and ϕ212-250 μm) of closed and open HA microspheres were created. Bone regeneration was conducted with a rat calvarial defect model. No osteoinductive agents were added in order to distinguish the intrinsic osteogenic properties of the open HA microspheres.

## Materials and Method

### Preparation of closed and open hollow hydroxyapatite (HA) microspheres

The closed hollow HA microspheres were prepared by conversion solid glass microspheres in aqueous phosphate solution as described in a previous study. Briefly, calcium-lithium-borate glass with the composition of 15CaO, 11Li_2_O and 74B_2_O_3_ (wt. %), designated as CaLB3-15, was prepared by melting CaCO_3_, Li_2_CO_3_, H_3_BO_3_ (Alfa Aesar, Haverhill, MA, USA) in a platinum crucible at 1200 °C for 45 min and then quenching the melt between stainless steel plates. Glass particles of were obtained by grinding the glass *via* a mortar and pestle, crashing in a shatter box and sieving through 100 and 140 mesh sieves for ϕ106-150 μm in size, or 60 and 70 mesh sieves for ϕ212-250 μm in size. Glass microspheres were obtained by dropping the crushed particles down through a vertical furnace at 1200 °C. The closed hollow hydroxyapatite microspheres were obtained by reacting the glass microspheres in a 0.02 M K_2_HPO_4_ solution at 37 °C and pH = 9 for 7 days. In the conversion process, 1 g glass was immersed in a 200 ml phosphate solution and the system was stirred gently and continuously. The converted microspheres were washed with distilled water and anhydrous ethanol, and then dried at room temperature for at least 12 h and at 90 °C for at least 12 h.

The open hollow HA microspheres were obtained by sectioning the closed hollow HA microspheres using a microtome. Briefly, the closed HA microspheres were fixed on a wax block using a water-soluble tape and were sectioned by microtome. The open HA microspheres were washed with distilled water and ethanol, and then dried at room temperature for at least 12 h and at 90 °C for at least 12 h. The debris in open HA microspheres were removed using sieves.

### Characterization of closed and open hollow hydroxyapatite (HA) microspheres

The microstructures of the closed HA microspheres, cross-section of closed HA microspheres, and open HA microspheres were observed using a scanning electron microscope (SEM; S4700 Hitachi, Tokyo, Japan) with an accelerating voltage of 15kV and working distance at 12 mm. The local composition of the surface layer, middle layer and inner layer of the mesoporous shell wall of the HA microspheres was investigated using energy dispersive X-ray (EDS) analysis in SEM with an electron beam spot size of 1 μm.

The specific surface area (SSA) of the closed and open HA microspheres and pore size distribution of the shell wall were measured by using nitrogen absorption (Autosorb-1; Quantachrome, Boynton Beach, FL) as described in a previous study. Three hundred milligrams of closed or open HA microspheres were weighted and evacuated at 120 °C for 15 h to remove absorbed moisture. The volume of nitrogen absorbed and desorbed at different relative gas pressure was measured and used to construct adsorption-desorption isotherms. The first twelve points of the adsorption isotherm, which initially followed a linear trend implying monolayer formation of adsorbate, were fitted to the Brunauer-Emmett-Teller equation to determine the specific surface area. The pore size distribution of the shell wall of the hollow HA microspheres was calculated using the Barrett-Joiner-Halenda method applied to the deposition isotherms [18].

### Animals and surgical procedures

All animal use and care procedures were approved by the Missouri S&T Institutional Animal Care and Use Committee in compliance with the NIH Guide for Care and Use of Laboratory Animals (1985). The rat calvarial defects were implanted with four groups of implants composed of closed or open hollow HA microspheres for 6 weeks and 12 weeks (Table 1). The implantation time was based upon considerable bone regeneration in rat calvarial defects implanted with hollow HA microspheres observed in previous studies. The closed or open HA microspheres of ϕ212-250 μm were randomly implanted to defect areas. The closed or open HA microspheres of ϕ106-150 μm microspheres were randomly implanted to defect areas, but mixing implants of closed and open microspheres in the same animal was avoided due to the possible migration of low-weight open HA microspheres.

**Table 1.** Implants groups composed of closed or open hollow hydroxyapatite microspheres.

| Group | HA microspheres |  | Sample size (n) |  |
| --- | --- | --- | --- | --- |
|  |  |  | 6 weeks | 12 weeks |
| 1 | 106-150 $\mu\text{m}$ | Closed | 5 | 5 |
| 2 |  | Open | 5 | 5 |
| 3 | 212-250 $\mu\text{m}$ | Closed | 5 | 10 |
| 4 |  | Open | 5 | 10 |

The male Sprague-Dawley rats (3 months old, weight = 350 ± 30 g, Envigo, USA) were acclimated for 2 weeks to diet, water, and housing under a 12 h/12 h light/dark cycle. The rats were anesthetized with a combination of ketamine and xylene (0.15 μl per 100 g) and maintained under anesthesia with isoflurane in oxygen. The surgery area was shaved, scrubbed with 70% ethanol and iodine, and draped. With sterile instruments and using an aseptic technique, a 1 cm cranial skin incision was made in an anterior to posterior direction along the midline. The subcutaneous tissue, musculature and periosteum were dissected and reflected to expose the calvaria. Bilateral full thickness defects (4.6 mm in diameter) were created in the central area of each parietal bone using a saline-cooled trephine drill. The sites were constantly irrigated with sterile PBS to prevent overheating of the bone margins and to remove the bone debris. Each defect was randomly implanted with HA microspheres of each group. After the implantation of the hollow HA microspheres, one drop of Ringer’s solution was added to each defect. The periosteum and skin were repositioned and closed with wound clips. Each animal received an intramuscular injection of ~200 μl buprenorphine and ~200 μl penicillin post-surgery. All animals were monitored daily for the condition of the surgical wound, food intake, activity and clinical signs of infection. After 6 weeks, the animals were sacrificed by CO_2_ inhalation, and the calvarial defect sites with surrounding bone and soft tissue were harvested for subsequent evaluations.

### Histological processing

Harvested calvarial samples were fixed in a 10% formaldehyde solution for five days. The samples were cut into half after being washed with deionized water. Half of the sample was for paraffin embedding, and the other half was for poly (methyl methacrylate) (PMMA) embedding. The paraffin-embedded samples were decalcified in 14 wt. % ethylenediaminetetraacetic acid (EDTA, Sigma-Aldrich, USA) for 2 weeks, dehydrated in ethanol, and then embedded in paraffin using standard histological techniques. These samples were sectioned using microtome. The thickness of the tissue section with paraffin was 5 μm. These slices were then stained with hematoxylin and eosin (H&E) [19]. Without decalcification, the samples for PMMA embedding were dehydrated in ethanol and embedded in PMMA. These samples were sectioned, affixed to acrylic slices, and ground to a thickness down to 50 μm using a micro-grinding system (EXAKT 400CS, Norderstedt, Germany). The von Kossa staining was used to observe mineralization [20].

### Histomorphometric analysis

Histomorphometric analysis was carried out using optical images of stained sections and Image J software (National Health Institute, USA). The percentage of new bone formed in calvarial defect was evaluated from the H&E stained sections. The newly formed bone was identified by outlining the edge of the defect, with the presence of old and new bone being identified by lamellar and woven bone, respectively. The total defect area was measured from one edge of the old calvarial bone, including the entire implant and tissue within it, to the other edge of the old bone. The newly formed bone within this area was then outlined and measured; the amount of the new bone was expressed as a percentage of the total defect area. The amount of von Kossa positive area was shown as a percent of the total defect area.

### Statistical analysis

Measurements of the percentage of new bone (relative to the entire defect area) were expressed as a mean ± SD. Analysis for differences between groups was performed using one-way analysis of variance (ANOVA) followed by the Tukey’s post hoc test; the differences were considered significant at P < 0.05.

## Results

### Geometry of the closed and open hydroxyapatite microspheres

The closed HA microspheres were prepared by converting glass microspheres in a phosphate solution. The diameters of the starting glass microspheres were ϕ106-150 μm and ϕ212-250 μm, respectively. After conversion, changes in the diameter of the microspheres were negligible. The SEM images revealed a spherical shape of closed HA microspheres with two size ranges: ϕ106-150 μm (thereafter, small size; Fig. 1A1 and A2) and ϕ212-250 μm (thereafter, large size; Fig. 1C1 and C2). Open HA microspheres were sectioned from closed HA microspheres using a microtone. The SEM images confirmed precise sectioning of open HA microspheres of both sizes (Fig. 1B1, B2, D1 and D2). Compared to the complete spherical structure of closed HA microspheres, the open HA microspheres were near hemispherical. The hollow microsphere had a mesoporous shell and a hollow core (0.6 of the microsphere diameter). The shell wall consisted of two distinct layers: a denser external layer and a more porous internal layer. For both size ranges of HA microspheres, the thickness of the denser layer was ~ 5 μm. The open HA microspheres of both sizes showed the dense external part and rough and porous internal part of the shell wall (Fig. 2). Both size ranges of the closed and open HA microspheres showed similar microstructures of the shell wall. The HA microspheres were formed by needle-like hydroxyapatite nanoparticles. The external surface tended to be denser than the internal surface.

**Figure 1.**
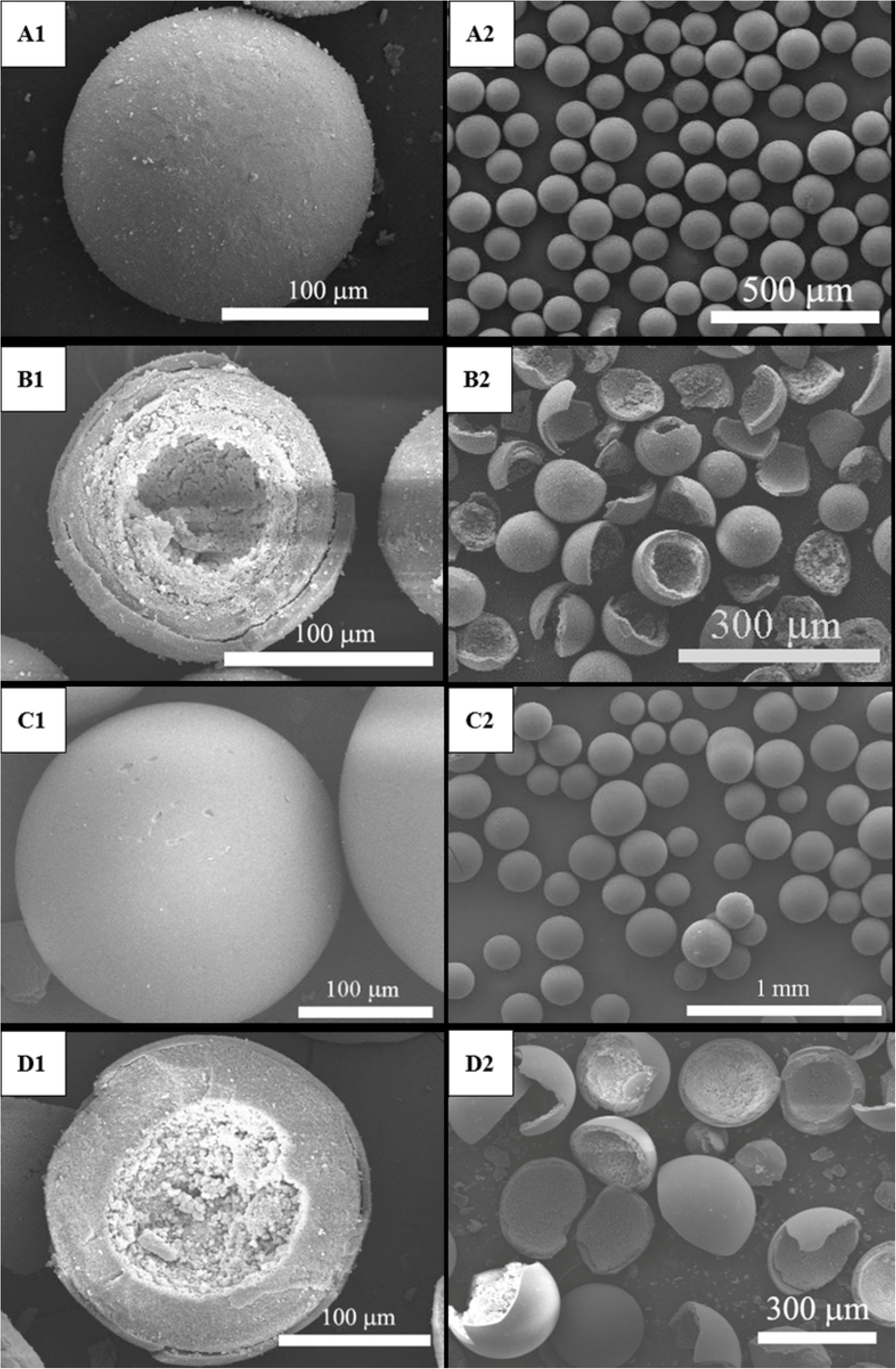
SEM images of 106-150 μm closed HA microspheres (A1, A2) and open HA microspheres (B1, B2) and 212-250 μm closed HA microspheres (C1, C2) and open HA microspheres (D1, D2).

**Figure 2.**
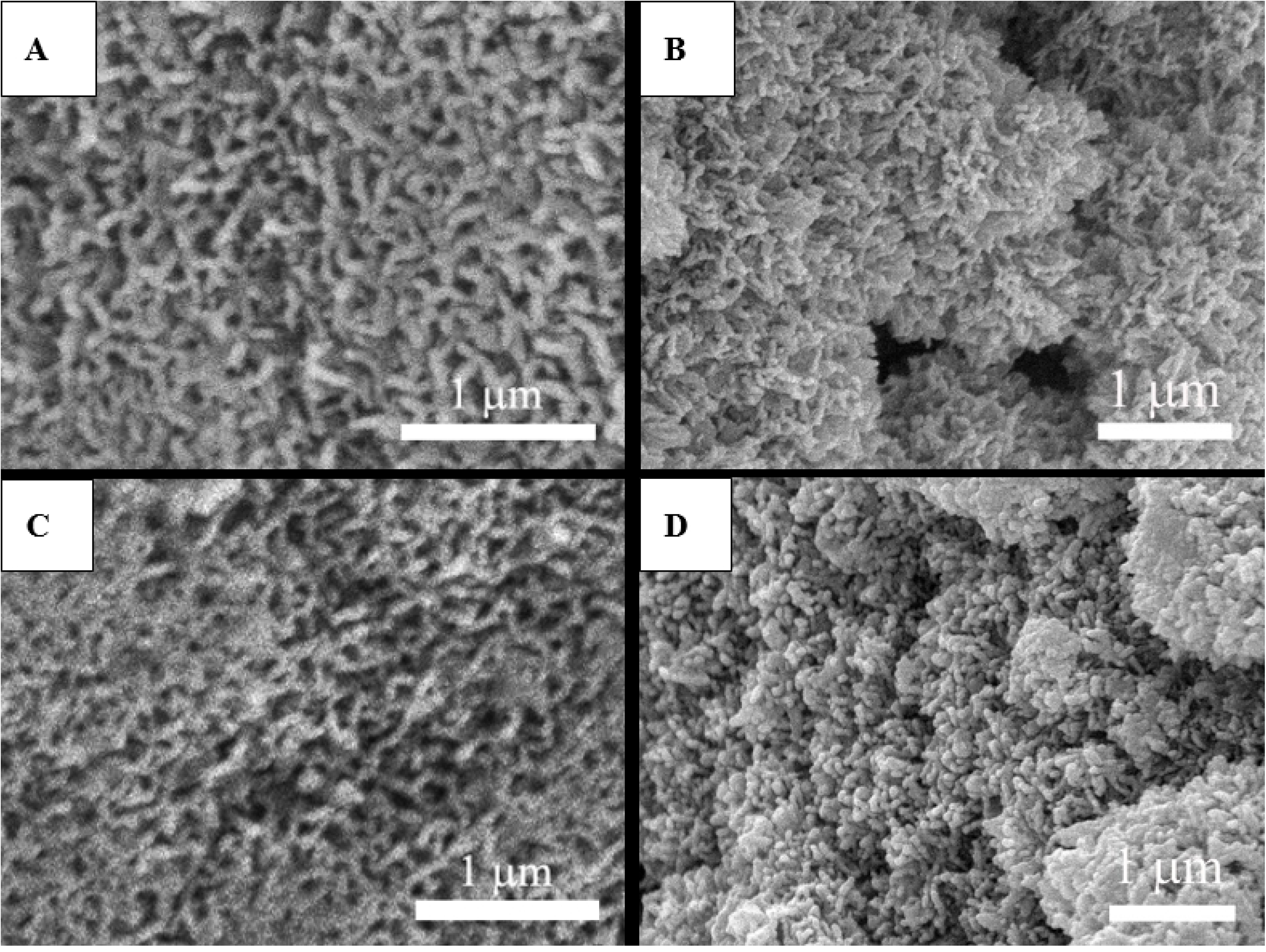
SEM images of external surface (A) and internal surface (B) of 106-150 μm open HA microspheres and external surface (C) and internal surface (D) of 212-250 μm open HA microspheres.

The BET surface area and average pore size of closed HA microspheres in two size ranges are summarized in Table 2. The surface areas of small and large closed HA microspheres were 101 m^2^/g and 168 m^2^/g, respectively. The average pore sizes of small and large closed HA microspheres were 13 nm and 10 nm, respectively. The surface area was higher in the large HA microspheres, while the average pore size was higher in the small HA microspheres.

**Table 2.**
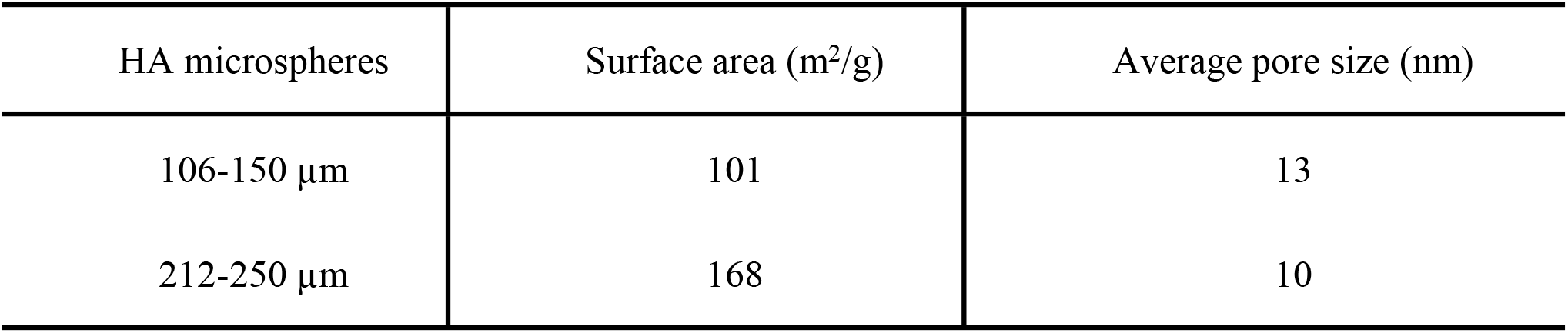
Surface area and average pore size of 106-150 μm and 212-250μm HA microspheres.

| HA microspheres | Surface area ($\text{m}^2/\text{g}$) | Average pore size (nm) |
| --- | --- | --- |
| 106-150 $\mu\text{m}$ | 101 | 13 |
| 212-250 $\mu\text{m}$ | 168 | 10 |

### Composition of the closed and open hollow hydroxyapatite microspheres

A high-resolution cross-section of the hollow HA microspheres in both sizes is shown in Fig. 3. The shell walls of the microspheres were divided into three regions: external layer, middle layer and inner layer. Compositions at the midpoint of each region were analyzed by EDS for the Ca/P atomic ratio (Table 3). The Ca/P atomic ratios of the HA microspheres of small size from the surface layer to the inner layer were 1.63 ± 0.11, 1.63 ± 0.11, and 1.60 ± 0.14. The Ca/P atomic ratios of the HA microsphere of large size from the surface layer to the inner layer were 1.67 ± 0.10, 1.63 ± 0.08 and 1.63 ± 0.06. There was no significant difference in Ca/P ratio within the three regions or between the two size ranges of HA microspheres (n=10, p>0.05). The Ca/P atomic ratios of the three regions were close to the theoretical Ca/P value of stoichiometric hydroxyapatite, 1.67.

**Table 3.** Ca/P atomic ratio (n = 10; mean ± SD) for the three regions for 106–150 µm and 212–250 µm.

| HA microspheres | Cross-sectional zone | Ca/P atomic ratio (mean ± SD) |
| --- | --- | --- |
| 106-150 $\mu\text{m}$ | Surface layer (A) | $1.63 \pm 0.11$ |
| | Middle layer (B) | $1.63 \pm 0.11$ |
| | Inner layer (C) | $1.60 \pm 0.14$ |
| 212-250 $\mu\text{m}$ | Surface layer (D) | $1.67 \pm 0.10$ |
| | Middle layer (E) | $1.63 \pm 0.08$ |
| | Inner layer (F) | $1.63 \pm 0.06$ |

**Figure 3.**
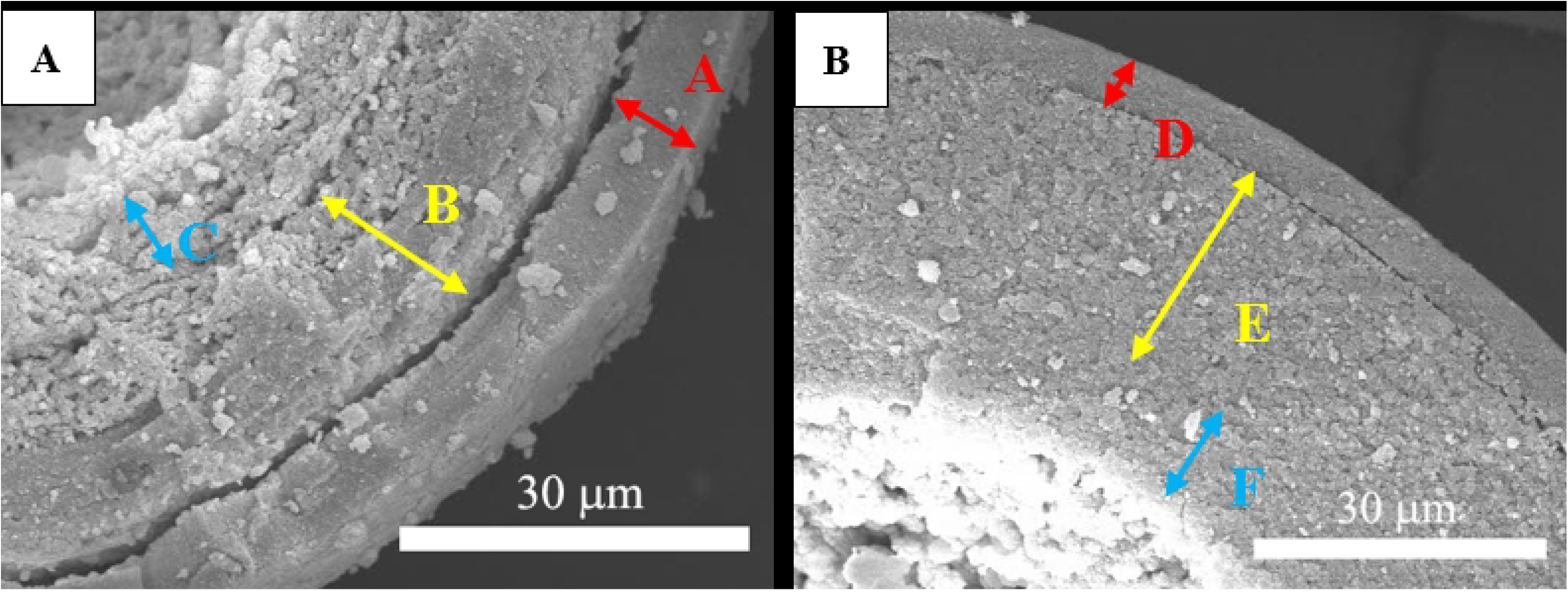
SEM images of cross section of 106-150 μm open HA microspheres (A) and 212-250μm open HA microspheres (B).

### Evaluation of bone regeneration in rat calvarial defects

H&E and von Kossa stained sections of the implants with closed and open hollow HA microspheres of the two size ranges after 6 weeks in rat calvarial defects are shown in Fig. 4 and Fig. 5. Bone regeneration was limited and confined mainly to the edge of the defects and some bone bridging along the bottom of implants. Fibrous tissues (light blue in H&E stained sections) filled the space between the microspheres. New bone formation in the implants with the smaller size of closed and open HA microspheres was 12 ± 3% and 17 ± 6%, respectively (Fig. 4 and Table 4). The von Kossa positive areas in in the implants with the smaller size of closed and open HA microspheres were 41 ± 3% and 49 ± 5%, respectively (Table 5). The percentages of new bone in the implants with the larger size of closed and open HA microspheres were 6 ± 2 % and 12 ± 3 %, respectively (Fig. 5). The von Kossa positive areas in the implants with the larger size of closed and open HA microspheres were 30 ± 3% and 35 ± 3%, respectively. Open HA microspheres showed significant improvement in bone regeneration compared with closed HA microspheres for both size ranges at 6 weeks in rat calvarial defects (n = 5, *p’s* < 0.05 for both sizes, Fig. 8 and 9). Smaller closed HA microspheres showed a significant increase in bone regeneration than the larger closed HA microspheres (n = 5, *p* < 0.05). Based on the H&E results, there was a borderline difference in new bone formation between the two size ranges of open HA microspheres (n = 5, p = 0.050). However, based on von Kossa results, the smaller open HA microspheres showed a significant enhancement in bone growth compared to the larger open HA microspheres (n = 5, *p* < 0.001).

**Table 4.** Comparative new bone formation in all implants after 6 or 12 weeks based on H&E staining. The amount of new bone is expressed as a percent of the total defect area (mean ± SD).

| Hollow HA microspheres |  | New bone (%) |  |
| --- | --- | --- | --- |
|  |  | 6 weeks | 12 weeks |
| φ106-150 μm | Closed | 12 ± 3 | 26 ± 8 |
|  | Open | 17 ± 6 | 49 ± 7 |
| φ212-250 μm | Closed | 6 ± 2 | 30 ± 9 |
|  | Open | 12 ± 3 | 40 ± 8 |

**Figure 4.**
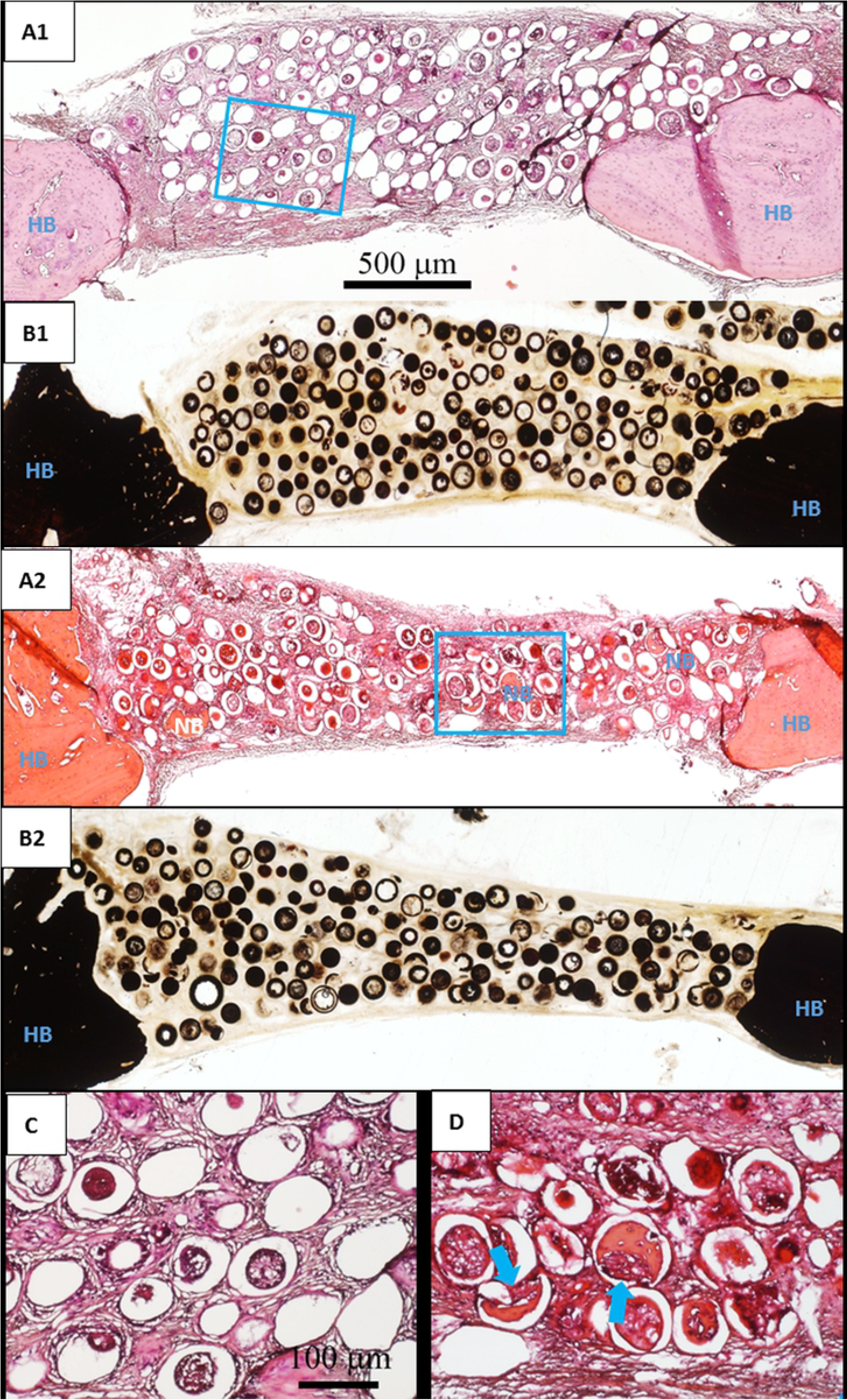
H&E stained and von Kossa sections of implants composed of closed (A1, B1) and open (A2, B2) hollow HA microspheres (ϕ106-150 μm) after 6 weeks in rat calvarial defects; (C, D) higher-magnification images of boxed area in (A1, A2). HB: host bone; NB: new bone. Blue arrow: new bone growth in micro-concavity.

**Figure 5.**
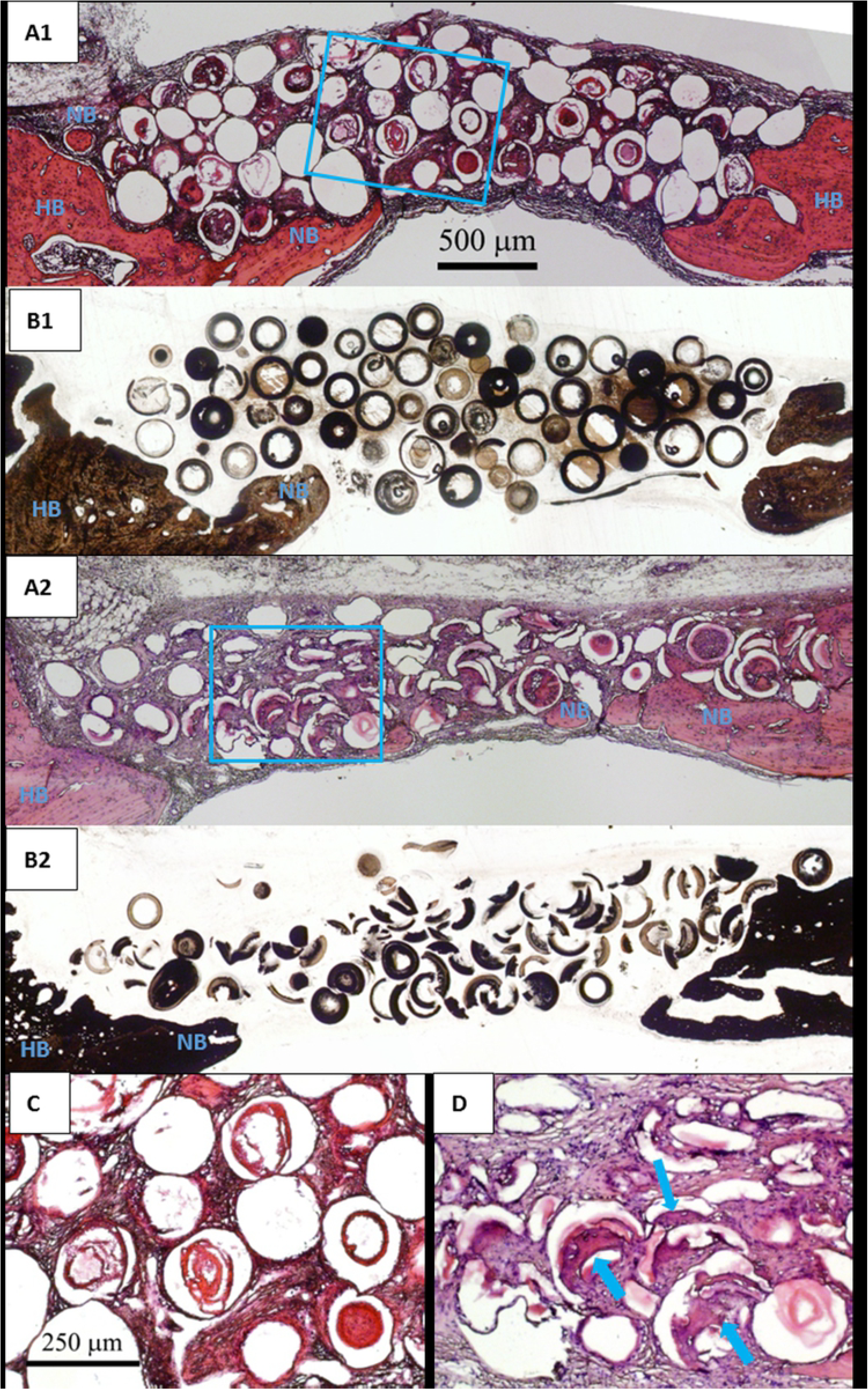
H&E and von Kossa stained sections of implants composed of closed (A1, B1) and open (A2, B2) HA microspheres (ϕ212-250 μm) after 6 weeks in rat calvarial defects; (C, D) higher-magnification images of boxed area in (A1, A2). HB: host bone; NB: new bone. Blue arrow: new bone growth in micro-concavity.

Higher magnification images of the closed and open HA microspheres of both sizes are shown in Fig. 4C and D (from the boxed areas of Fig. 4A1 and A2) and Fig. 5C and D (from the boxed areas in Fig. 5A1 and A2). For the closed HA microspheres in both size ranges, bone formation was scanty, while the fibrous tissues filled the pore space between the closed HA microspheres and infiltrated into the hollow core of some broken closed HA microspheres. In comparison, more bone regeneration was observed in the micro-concavity of open HA microspheres (indicated by blue arrows) in both sizes (ϕ106-150 μm and ϕ212-250 μm).

The outcomes from the implants with the closed and open HA microspheres of the two size ranges after 12 weeks in rat calvarial defects are shown in Figs. 6 and 7. New bones were formed from the edge of the defects and on the bottom of the implants. For the open HA microspheres of both size ranges, more new bone growth in the micro-concavity can be found; the remaining open HA microspheres can be observed in the new bone bridging the ends of defects. For the closed and open HA microspheres of small size (Fig. 6), the percentages of new bone formation were 26 ± 8% and 49 ± 7%, respectively; the von Kossa positive areas were 55 ± 5% and 76 ± 4%, respectively. For the closed and open HA microspheres of large size (Fig. 7), the percentages of new bone were 30 ± 9% and 40 ± 8 %, respectively; the von Kossa positive areas were 56 ± 5% and 65 ± 5%, respectively. The open HA microspheres showed significant improvement in bone regeneration when compared to the closed HA microspheres in both size ranges during a period of 12 weeks in rat calvarial defects (n = 5, *p’s* < 0.001 for small size; n = 5, *p’s* < 0.05 for large size). There was no significant difference in new bone formation between the two size ranges of closed HA microspheres (n = 5~10, *p* > 0.05). However, smaller open HA microspheres showed a more significant increase in bone regeneration than larger closed HA microspheres (n = 5, *p* < 0.05). Bone regeneration was time-dependent for both size ranges; new bone formation increased significantly from 6 weeks to 12 weeks in rat calvarial defects (n = 5, *p’s* < 0.001 for closed HA microspheres; n = 5~10, *p’s* < 0.001 for open HA microspheres).

**Figure 6.**
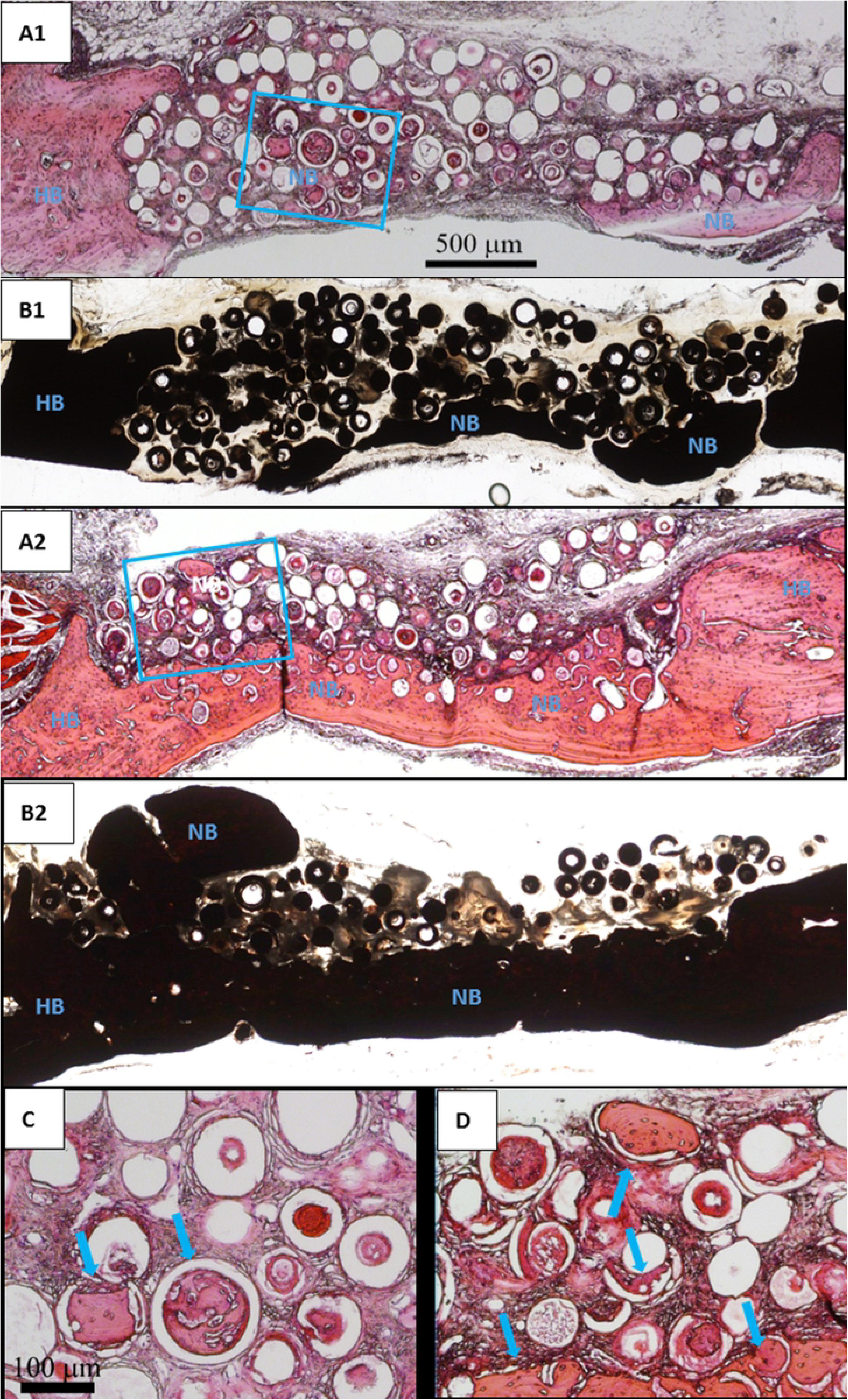
H&E and von Kossa stained sections of implants composed of closed (A1, B1) and open (A2, B2) HA microspheres (ϕ106-150 μm) after 12 weeks in rat calvarial defects; (C, D) higher-magnification images of boxed area in (A1, A2). HB: host bone; NB: new bone. Blue arrow: new bone growth in micro-concavity.

**Figure 7.**
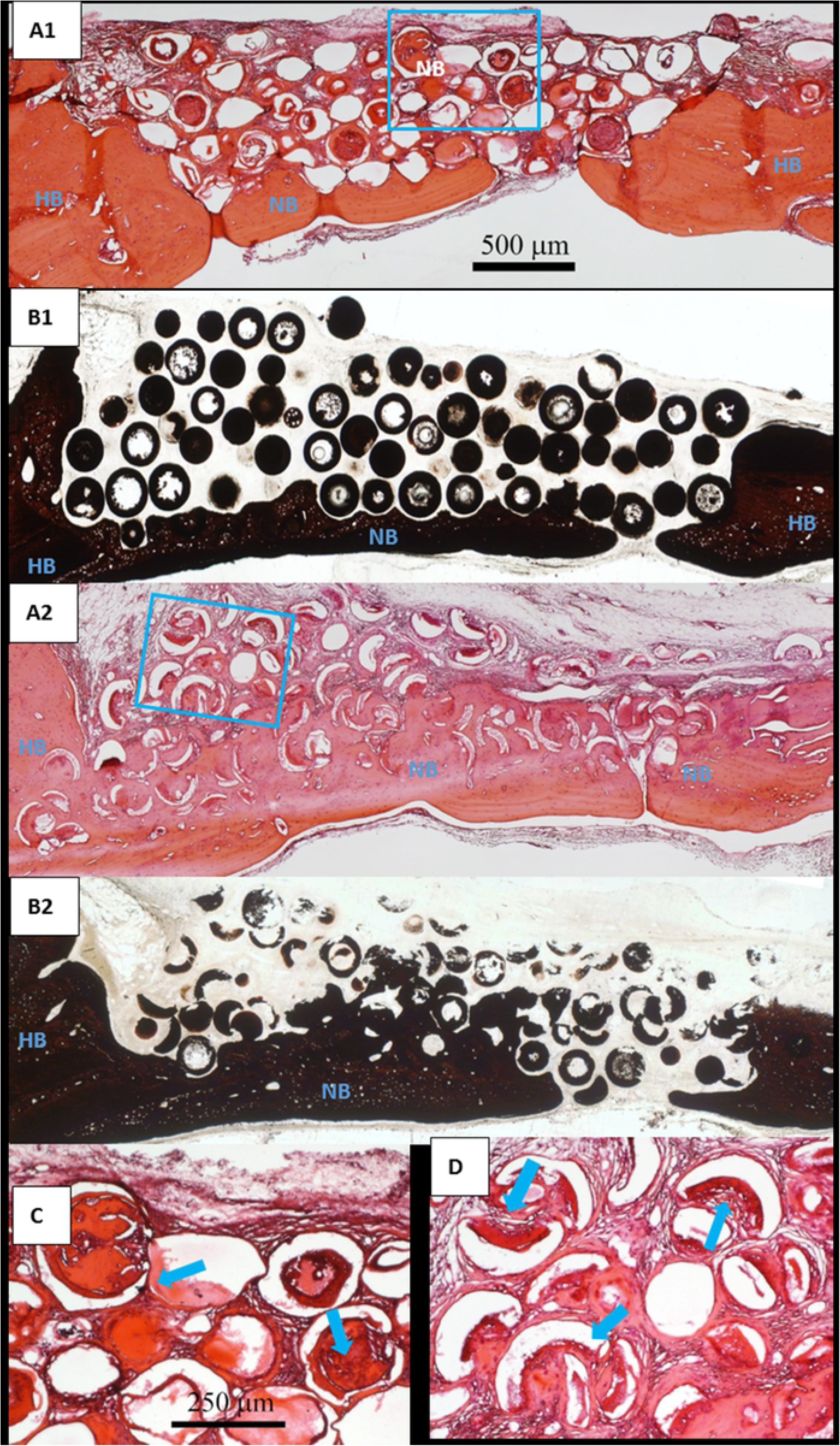
H&E and von Kossa stained sections of implants composed of closed (A1, B1) and open (A2, B2) HA microspheres (ϕ212-250 μm) after 12 weeks in rat calvarial defects; (C, D) higher-magnification images of boxed area in (A1, A2). HB: host bone; NB: new bone. Blue arrow: new bone growth in micro-concavity.

A comparison of closed and open HA microspheres in both sizes at 12 weeks is shown in higher magnified images in Fig. 6C and D (from the boxed areas of Fig. 6A1 and A2) and Fig. 7C and D (from the boxed areas of Fig. 7A1 and A2). Bone regeneration in the cores of some broken closed HA microspheres was identified. A higher degree of new bone formation in the micro-concavity of open HA microspheres was observed.

**Figure 8.**
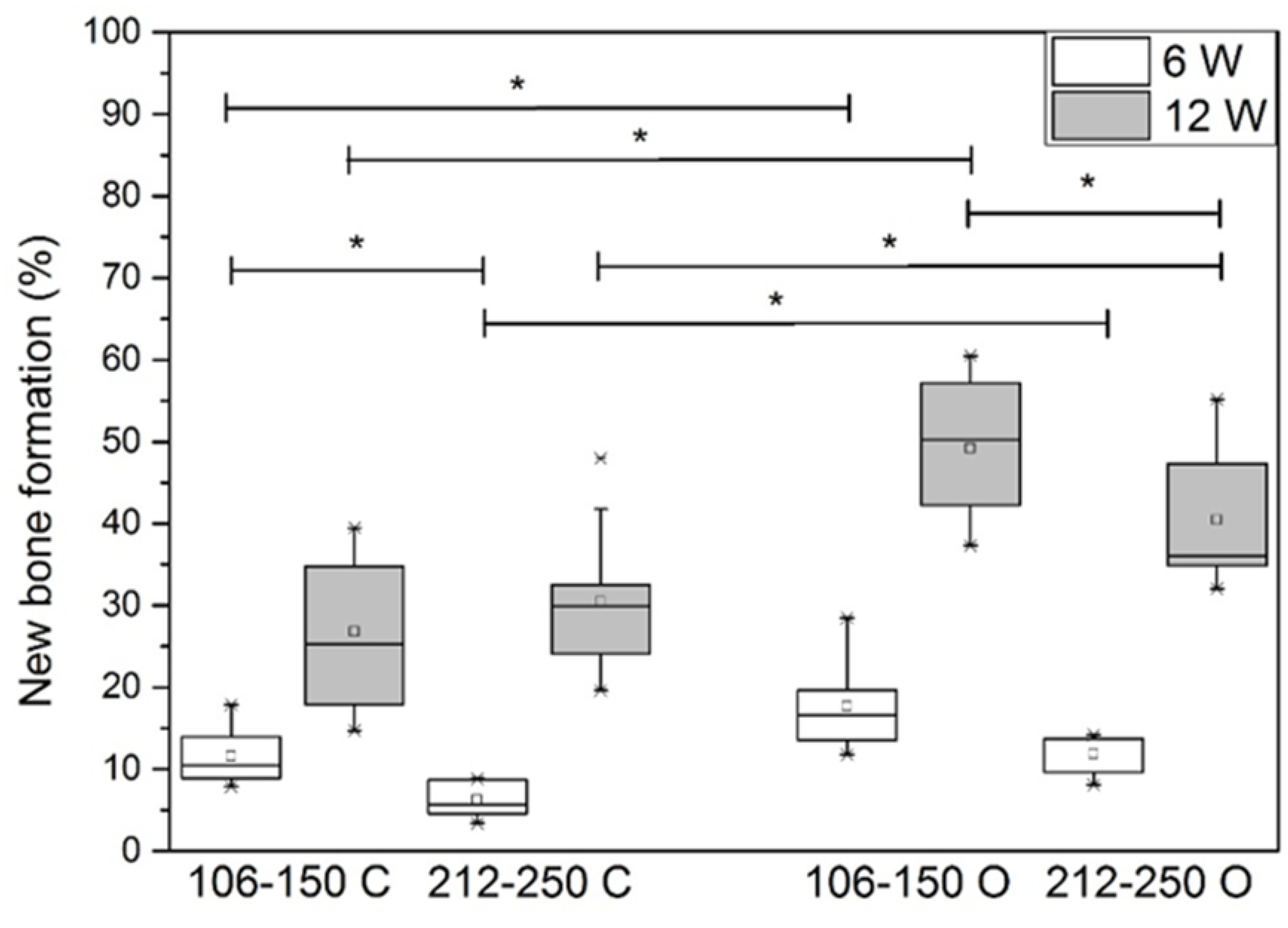
Comparative new bone formation in implants with closed and open hollow HA microspheres with diameter of 106-150 μm or 212-250 μm after 6 weeks (6 W) and 12 weeks (12 W) in rat calvarial defects (Mean ± SD; n = 5~10, * significant difference between groups; *p* < 0.05).

**Figure 9.**
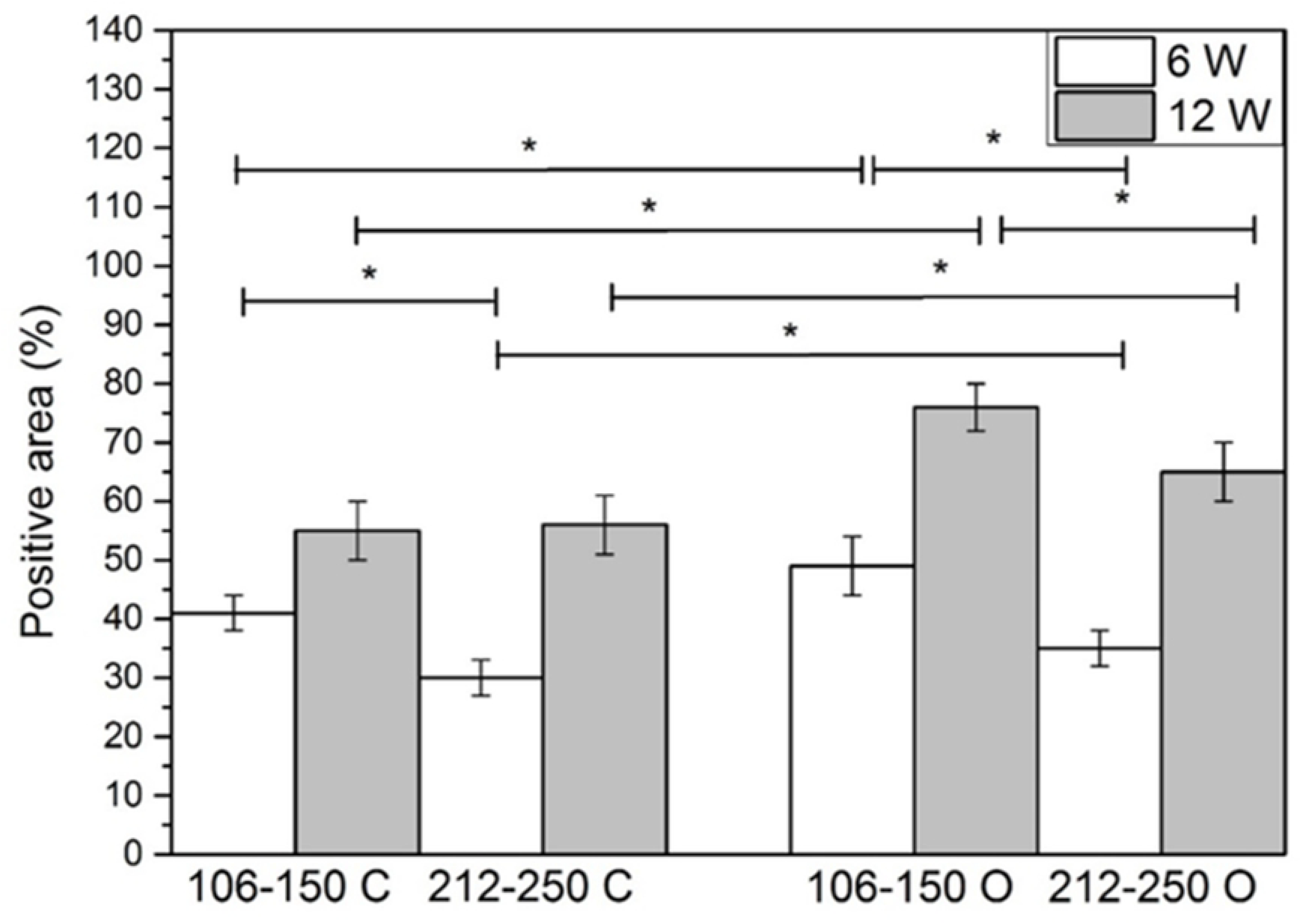
Comparative von Kossa positive area for implants of closed and open hollow HA microspheres with diameter of 106-150 μm or 212-250 μm after 6 weeks (6 W) and 12 weeks (12 W) in rat calvarial defects (Mean ± SD; n = 5~10, * significant difference between groups; *p* < 0.05).

## Discussion

The capability of HA microspheres to regenerate bone can presumably be affected by the differences between closed and open HA microspheres in microstructure. In this study, the microstructure of closed and open HA microspheres in two size ranges (ϕ106-150 μm vs. ϕ212-250 μm) were analyzed. To test HA microspheres in facilitating bone regeneration, rat calvarial defects were created and HA microspheres were implanted. Bone regeneration was evaluated in weeks 6 & 12.

For both size ranges, the thickness of the denser (outer) layer was ~ 5 μm, while the ratio of the hollow core diameter to the microsphere diameter is ~0.6. The factors leading to these two distinct layers are still unclear. In the glass conversion process [16, 21-23], ions are dissolved from glass (i.e., Ca^2+^, Li^+^, B^3+^) to the aqueous solution. The Ca^2+^ from glass reacts immediately with phosphate anions from solution to form calcium phosphate. The calcium phosphate precipitates onto the glass surface due to its insolubility in the system. As the glass dissolves, the calcium phosphate layer continues to thicken until the glass is completely converted to calcium phosphate. The kinetics and mechanism of the formation of the HA layer in borate glass is investigated in several studies [22, 24-26]. The conversion rate is initially described by a reaction-controlled model (linear kinetics); however, at the later stage, a three-dimensional diffusion model (parabolic kinetics) better explains the conversion rate. Presumably, the denser layer and porous layer results from these two kinetic models. Additional experiments can be set-up to further investigate the dynamic changes of SSA and pore size.

Our *in vivo* experiment showed the effectiveness of open HA microspheres in bone regeneration. For both size ranges of the open HA microspheres, new bone formation was observed in both 6 weeks and 12 weeks post-implantation. The amount of new bone growth increased from 6 weeks to 12 weeks. In the study of 12-week implantation with small microspheres, new bone formation with the implants of open microsphere was about twice that of the closed microspheres; for large microspheres, new bone formation in the implants of open microspheres was about 30% higher than that of the closed microspheres. Thus, the open microspheres were more effective in facilitating bone regeneration than the closed microspheres. Compared to the closed microsphere, the open microsphere had a micro-concave region with a more porous and rougher surface (see Fig 1&2). These characters (i.e., micro-concavity, porosity, roughness) could contribute to the difference in bone regeneration between the closed and open microspheres.

The effectiveness of micro-concavity in bone regeneration has been investigated by others [27–32]. Substantial mineralization of simulated body fluid on the discs made of calcium phosphate ceramic were observed inside concavities but not at the planar surface [31]. Smaller concavity (0.4 mm in diameter) can induce much more mineralization than larger concavities (0.8 mm or 1.8 mm in diameter) [31]. An *in vivo* study demonstrated that concavity appeared to stimulate formation of blood vessels, a critical process for bone formation [32]. Stem cells showed better outcomes on a concave surface than a flat surface in terms of cell maturation, osteodifferentiation, and specific protein production [28]. Bone formation by intramembranous ossification preferred to occur on a concave surface as well [30]. Concavity is also conducive to accumulation of growth factors such as BMPs [27]. Differences in microstructure may also be a contributing factor to the outcome of bone regeneration. The internal concave surface was more porous and rougher compared to external convex surface.

The differences in porosity and roughness could influence dissolution/degradation of biomaterials, adsorption of growth factors, and mineral deposition from body fluid [33–40]. For instance, the degradation of a porous surface could lead to faster Ca^2+^ release which is a key factor in facilitating angiogenesis [41]. Further, a more porous and rougher surface could be a more suitable substrate for adsorption of biologically active molecules, such as BMPs and growth factors. Together, these lead to enhanced cell attachment, proliferation and differentiation.

Dissolution/degradation of HA have been shown to be affected by the ratio of Ca/P of the microspheres [42–44]. The dissolution of HA in water increased as the Ca/P ratio decreased [42]. Higher dissolution/degradation of HA could release more Ca^2+^ and phosphate ions, which could facilitate bone regeneration. In this study, there was no significant difference in Ca/P ratio within the three regions or between the two size ranges of HA microspheres. It is possible that the Ca/P ratio of our HA microspheres can be manipulated to achieve varying degree of dissolution.

The current study demonstrated that the small open microspheres induced a more significant increase in bone regeneration than the large open microspheres at both 6 weeks and 12 weeks. One reason for this difference may be attributed to total surface area on microspheres where cells can be attached to. A simulation of the difference in available surface area for cell attachment was made. Given the same mass, the same size distribution pattern of the open and closed microspheres of the same size, and the same density of the shell of the microspheres of different sizes, the open microspheres of the same size have larger surface area than that of the closed microspheres for cells to attached to. For instance, the closed and open microspheres of ϕ106-150 μm have surface areas of 584 cm^2^ (assuming the total volume of the microsphere shell is 1 cm^3^) and 981 cm^2^, respectively. The closed and open microspheres of ϕ212-250 μm surface areas of have 328 cm^2^ and 552 cm^2^, respectively. Another reason for the difference in bone regeneration could be due to the curvature. The small microspheres have higher curvature than the large microspheres. It remains to be investigated how the curvature of the microspheres affect cellular physiology leading to the differential outcome of bone regeneration.

An apparent observation is that new bone formation with implants of the small open microspheres was able to completely bridge the defects at the bottoms of all the implants. In comparison, not all animals with closed microspheres were able to bridge the entire defects. During the regeneration process, new bone formation started from the edge of the host bone and from the bottom of the defect (dura matter), where osteogenic cells and blood supply were abundant. The open microspheres might absorb the osteogenic factors by diffusion or fluid transport and trigger bone growth in the micro-concavity. The open microspheres at the bottom of the implants had the best chance of contact with the osteogenic factors not only from dura matter but also from the edges induced by the open microsphere in periphery. We observed that a large number of smaller pieces of open microspheres was found in the bottom of the implants. This might be caused by the rats’ physical activity of daily living.

In this work, the closed HA microspheres of ϕ106-150 µm significantly enhance bone regeneration than those of ϕ 212-250 µm at 6 weeks; no significant difference in bone regeneration between two size ranges at 12 weeks. Compared to the work by Fu [14], new bone formation with the closed HA microspheres of ϕ150-250 µm was significantly greater than that with the closed HA microspheres of ϕ106-150 µm at 12 weeks. It should be noted that there is a significant difference in the size range of the large microspheres between these two studies; thus, they should not be viewed as conflicting results.

## Conclusion

The open HA microspheres significantly enhance the bone regeneration as compared to the closed HA microspheres at both 6 weeks and 12 weeks. Compared with the larger size of open HA microspheres (smaller curvature), the smaller size of open HA microspheres (larger curvature) resulted in a more significant increase in bone regeneration. The differences in microstructures of the HA microspheres (i.e., curvature, concavity, porosity, surface roughness, total surface area available for cells to attached to) may deserve future attention of investigation.

## Author Contributions

Conceived and designed the experiments: MR YL YWH. Performed the experiments: YS. Analyzed the data: YS MR YL YWH. Wrote the paper: YS MR YWH.

## Acknowledgements

The work is financially supported by National Institute of Health [Grant #1R15DE023987-01]. We thank technical support by the S&T Material Research Center and the Department of Biological Sciences.

## Conflicts of Interest

The authors declare that there are no conflicts of interest.

## Abbreviations

BSA: bovine serum albumin
HA: hydroxyapatite
PBS: phosphate Buffer Saline
BMP-2: bone morphogenetic protein-2
H&E: hematoxylin and eosin
FBS: fetal bovine serum
EDTA: ethylenediaminetetraacetic acid

## References

1. Campana V, Milano G, Pagano E, Barba M, Cicione C, Salonna G, et al. Bone substitutes in orthopaedic surgery: from basic science to clinical practice. Journal of Materials Science: Materials in Medicine. 2014;25(10):2445–61. http://doi.org/10.1007/s10856-014-5240-2. PubMed PMID:24865980; PubMed Central PMCID: PMCPMC4169585.

2. Deschaseaux F, Sensébé L, Heymann D. Mechanisms of bone repair and regeneration. Trends in Molecular Medicine. 2009;15(9):417–29. http://doi.org/10.1016/j.molmed.2009.07.002. PubMed PMID: 19740701.

3. Loi F, Córdova LA, Pajarinen J, Lin T-h, Yao Z, Goodman SB. Inflammation, fracture and bone repair. Bone. 2015;86:119–30. http://doi.org/10.1016/j.bone.2016.02.020. PubMed PMID: 26946132; PubMed Central PMCID: PMCPMC4833637.

4. Kalfas IH. Principles of bone healing. Neurosurg Focus. 2001;10(4):E1. http://doi.org/10.3171/foc.2001.10.4.2. PubMed PMID: 16732625.

5. Scarano A, Degidi M, Iezzi G, Pecora G, Piattelli M, Orsini G, et al. Maxillary sinus augmentation with different biomaterials: a comparative histologic and histomorphometric study in man. Implant dentistry. 2006;15(2):197–207. http://doi.org/10.1097/01.id.0000220120.54308.f3. PubMed PMID: 16766904.

6. Trombelli L. Which reconstructive procedures are effective for treating the periodontal intraosseous defect? Periodontology 2000. 2005;37(1):88–105. http://doi.org/10.1111/j.1600-0757.2004.03798.x. PubMed PMID: 15655027.

7. Kao RT, Conte G, Nishimine D, Dault S. Tissue engineering for periodontal regeneration. Journal of the California Dental Association. 2005;33(3):205–15. http://doi.org/10.4103/1735-3327.107570. PubMed PMID: 15918402.

8. Pandit N, Malik R, Philips D. Tissue engineering: A new vista in periodontal regeneration. Journal of Indian Society of Periodontology. 2011;15(4):328–37. http://doi.org/10.4103/0972-124X.92564. PubMed PMID: 22368355; PubMed Central PMCID: PMCPMC3283928.

9. Dahlin C, Johansson A, Sahlgrenska a, Institute of Clinical Sciences SfAB, Orthopaedics DoB, Göteborgs u, et al. Iliac crest autogenous bone graft versus alloplastic graft and guided bone regeneration in the reconstruction of atrophic maxillae: a 5 - year retrospective study on cost - effectiveness and clinical outcome. Clinical Implant Dentistry and Related Research. 2011;13(4):305–10. http://doi.org/10.1111/j.1708-8208.2009.00221.x. PubMed PMID: 21087398.

10. Giannoudis PV, Dinopoulos H, Tsiridis E. Bone substitutes: an update. Injury. 2005;36(3):S20–S7. http://doi.org/10.1016/j.injury.2005.07.029. PubMed PMID: 16188545.

11. McAuliffe JA. Bone graft substitutes. Journal of Hand Therapy. 2003;16(2):180–7. http://doi.org/10.1016/S0894-1130(03)80013-3. PubMed PMID: 16359252.

12. Reikerås O, Shegarfi H, Naper C, Reinholt FP, Rolstad B. Impact of MHC mismatch and freezing on bone graft incorporation: An experimental study in rats. Journal of Orthopaedic Research. 2008;26(7):925–31. http://doi.org/10.1002/jor.20595. PubMed PMID: 18302282.

13. Von Recum A, Jacobi JE. Handbook of biomaterials evaluation: scientific, technical, and clinical testing of implant materials. 2nd ed. Philadelphia, PA: Taylor & Francis; 1999.

14. Fu H, Rahaman MN, Brown RF, Day DE. Evaluation of bone regeneration in implants composed of hollow HA microspheres loaded with transforming growth factor β1 in a rat calvarial defect model. Acta Biomaterialia. 2013;9(3):5718–27. http://doi.org/10.1016/j.actbio.2012.11.017. PubMed PMID: 23168225; PubMed Central PMCID: PMCPMC3562431.

15. Xiao W, Fu H, Rahaman MN, Liu Y, Bal BS. Hollow hydroxyapatite microspheres: A novel bioactive and osteoconductive carrier for controlled release of bone morphogenetic protein-2 in bone regeneration. Acta Biomaterialia. 2013;9(9):8374–83. http://doi.org/10.1016/j.actbio.2013.05.029. PubMed PMID: 23747325; PubMed Central PMCID: PMCPMC3732511

16. Fu H. Hollow hydroxyapatite microspheres as devices for controlled delivery of proteins and as scaffolds for tissue engineering: ProQuest Dissertations Publishing; 2012.

17. Rahaman MN, Xiao W, Liu Y, Bal BS, editors. Osteoconductive and osteoinductive implants composed of hollow hydroxyapatite microspheres2014.

18. Barrett EP, Barrett EP, Joyner LG, Joyner LG, Halenda PP, Halenda PP. The determination of pore volume and area distributions in porous substances. I. Computations from nitrogen isotherms. Journal of the American Chemical Society. 1951;73(1):373–80. http://doi.org/10.1021/ja01145a126.

19. Feng JQ, Zhang J, Dallas SL, Lu Y, Chen S, Tan X, et al. Dentin matrix protein 1, a target molecule for cbfa1 in bone, ss a unique bone marker gene. Journal of Bone and Mineral Research. 2002;17(10):1822–31. http://doi.org/10.1359/jbmr.2002.17.10.1822. PubMed PMID: 12369786.

20. Bonewald LF, Harris SE, Rosser J, Dallas MR, Dallas SL, Camacho NP, et al. Von Kossa staining alone is not sufficient to confirm that mineralization in vitro represents bone formation. Calcified Tissue International. 2003;72(5):537–47. http://doi.org/10.1007/s00223-002-1057-y. PubMed PMID: 12724828.

21. Fears KP. Formation of hollow hydroxyapatite microspheres 2001.

22. Nancollas GH, Zhang J. Formation and dissolution mechanisms of calcium phosphates in aqueous systems. In: Brown PW, Constantz B, editors. Hydroxyapatite and related materials. United State of America: CPC Press, Inc.; 1994. p. 73–81.

23. Wang Q, Huang W, Wang D, Darvell BW, Day DE, Rahaman MN. Preparation of hollow hydroxyapatite microspheres. Journal of Materials Science: Materials in Medicine. 2006;17(7):641–6. http://doi.org/10.1007/s10856-006-9227-5. PubMed PMID: 16770549.

24. Gu Y, Xiao W, Lu L, Huang W, Rahaman MN, Wang D. Kinetics and mechanisms of converting bioactive borate glasses to hydroxyapatite in aqueous phosphate solution. Journal of Materials Science. 2011;46(1):47–54. http://doi.org/10.1007/s10853-010-4792-x.

25. Huang W, Rahaman MN, Day DE, Li Y. Mechanisms for converting bioactive silicate, borate, and borosilicate glasses to hydroxyapatite in dilute phosphate solution. Physics and Chemistry of Glasses-European Journal of Glass Science and Technology Part B. 2006;47(6):647–58.

26. Yao A-H, Lin J, Duan X, Huang W-H. Formation mechanism of multilayered structure on surface of bioactive borosilicate glass. Chinese Journal of Inorganic Chemistry. 2008;24(7):1132–6.

27. Ripamonti U. Functionalized Surface Geometries Induce:“Bone: Formation by Autoinduction”. Frontiers in physiology. 2018;8:1084. http://doi.org/10.3389/fphys.2017.01084. PubMed PMID: 29467661; PubMed Central PMCID: PMCPMC5808255

28. Graziano A, d’Aquino R, Cusella-De Angelis MG, Laino G, Piattelli A, Pacifici M, et al. Concave Pit-Containing Scaffold Surfaces Improve Stem Cell-Derived Osteoblast Performance and Lead to Significant Bone Tissue Formation. PLoS ONE. 2007;2(6):e496. http://doi.org/10.1371/journal.pone.0000496. PubMed PMID: 17551577; PubMed Central PMCID: PMCPMC1876259

29. Gray C, Boyde A, Jones S. Topographically induced bone formation in vitro: implications for bone implants and bone grafts. Bone. 1996;18(2):115–23. http://doi.org/10.1016/8756-3282(95)00456-4. PubMed PMID: 8833205.

30. Coathup MJ, Hing KA, Samizadeh S, Chan O, Fang YS, Campion C, et al. Effect of increased strut porosity of calcium phosphate bone graft substitute biomaterials on osteoinduction. Journal of Biomedical Materials Research Part A. 2012;100A(6):1550–5. http://doi.org/10.1002/jbm.a.34094. PubMed PMID: 22419568.

31. Bianchi M, Urquia Edreira ER, Wolke JGC, Birgani ZT, Habibovic P, Jansen JA, et al. Substrate geometry directs the in vitro mineralization of calcium phosphate ceramics. Acta Biomaterialia. 2014;10(2):661–9. http://doi.org/10.1016/j.actbio.2013.10.026. PubMed PMID: 24184857.

32. Scarano A, Perrotti V, Artese L, Degidi M, Degidi D, Piattelli A, et al. Blood vessels are concentrated within the implant surface concavities: a histologic study in rabbit tibia. Odontology. 2014;102(2):259–66. http://doi.org/10.1007/s10266-013-0116-3. PubMed PMID: 23783569.

33. Yuan H, Yang Z, Li Y, Zhang X, De Bruijn JD, De Groot K. Osteoinduction by calcium phosphate biomaterials. Journal of Materials Science: Materials in Medicine. 1998;9(12):723–6. http://doi.org/10.1023/A:1008950902047. PubMed PMID: 15348929.

34. Danoux CB, Barbieri D, Yuan H, de Bruijn JD, van Blitterswijk CA, Habibovic P. In vitro and in vivo bioactivity assessment of a polylactic acid/hydroxyapatite composite for bone regeneration. Biomatter. 2014;4(1):e27664. http://doi.org/10.4161/biom.27664. PubMed PMID: 24441389; PubMed Central PMCID: PMCPMC3961484.

35. Fujibayashi S, Neo M, Kim H-M, Kokubo T, Nakamura T. Osteoinduction of porous bioactive titanium metal. Biomaterials. 2004;25(3):443–50. http://doi.org/10.1016/S0142-9612(03)00551-9. PubMed PMID: 14585692.

36. García-Gareta E, Hua J, Knowles JC, Blunn GW. Comparison of mesenchymal stem cell proliferation and differentiation between biomimetic and electrochemical coatings on different topographic surfaces. Journal of Materials Science: Materials in Medicine. 2013;24(1):199–210. http://doi.org/10.1007/s10856-012-4789-x. PubMed PMID: 23053816.

37. Barbieri D, Renard AJS, de Bruijn JD, Yuan H. Heterotopic bone formation by nano-apatite containing poly (D,L-lactide) composites. European cells & materials. 2010;19:252–61. http://doi.org/10.22203/eCM.v019a24. PubMed PMID: 20526989.

38. Kilpadi KL, Chang P-L, Bellis SL. Hydroxylapatite binds more serum proteins, purified integrins, and osteoblast precursor cells than titanium or steel. Journal of Biomedical Materials Research. 2001;57(2):258–67. http://doi.org/10.1002/1097-4636(200111)57:2<258::AID-JBM1166>3.0.CO;2-R. PubMed PMID: 11484189.

39. Bagambisa FB, Joos U. Preliminary studies on the phenomenological behaviour of osteoblasts cultured on hydroxyapatite ceramics. Biomaterials. 1990;11(1):50–6. http://doi.org/10.1016/0142-9612(90)90052-R. PubMed PMID: 2154267.

40. Hing KA. Bioceramic bone graft substitutes: Influence of porosity and chemistry. International Journal of Applied Ceramic Technology. 2005;2(3):184–99. http://doi.org/10.1111/j.1744-7402.2005.02020.x.

41. Aguirre A, González A, Navarro M, Castaño Ó, Planell JA, Engel E. Control of microenvironmental cues with a smart biomaterial composite promotes endothelial progenitor cell angiogenesis. European Cells and Materials. 2012;24:90–106. http://doi.org/10.22203/eCM.v024a07.

42. Mavropoulos E, Rossi AM, da Rocha NCC, Soares GA, Moreira JC, Moure GT. Dissolution of calcium-deficient hydroxyapatite synthesized at different conditions. Materials Characterization. 2003;50(2):203–7. http://doi.org/10.1016/S1044-5803(03)00093-7.

43. Driessens F. Formation and stability of calcium phosphates in relation to the phase composition of the mineral in calcified tissues. Bioceramics Calcium Phosphate: CRC Press; 2018. p. 1–32.

44. Tofighi A, Schaffer K, Palazzolo R, editors. Calcium phosphate cement (CPC): a critical development path. Key Engineering Materials; 2008: Trans Tech Publ.

